# Genomic plasticity of the *Azospirillum* genus in a biotechnological context

**DOI:** 10.1101/2025.10.16.682878

**Authors:** Sarah Henaut-Jacobs, Felipe F. Rimes-Casais, Gabriel Quintanilha-Peixoto, Gabriela Petroceli-Mota, Bruno da Costa Rodrigues, Isabella de Oliveira Pinheiro, Rodrigo Nunes-da-Fonseca, Fabio Lopes Olivares, Rampal S. Etienne, Thiago M. Venancio

## Abstract

Extensive agriculture and the use of chemical fertilizers cause notable environmental impacts on multiple levels, from reducing soil microbiota diversity to groundwater contamination. In this context, the usage of plant growth-promoting bacteria (PGPB) presents a sustainable alternative to enhance crop production while mitigating these adverse effects. *Azospirillum*, a bacterial genus renowned for its beneficial capabilities, particularly phytohormone production, is a key component of many commercial inoculants. In this work, we performed a comparative genomic analysis of all publicly available *Azospirillum* genomes and four novel isolates belonging to our microbial collection. Our analysis identified a species complex within the genus, which we designate the *A. brasilense* species complex, comprising species already used in commercial bioconsortia. This complex is characterized by a core set of exclusive genes linked to chemotaxis and host-recognition capability. Furthermore, we also validated the biosafety of the *A. brasilense* species complex and confirmed the plant growth-promoting potential of our novel isolates, highlighting their suitability for developing new biofertilizers.

## Introduction

Global agriculture faces the pressing challenge of ensuring food security without harming the environment and human health (FAO, 2017). Although critical since the Green Revolution, the historical over-reliance on chemical fertilizers and pesticides has led to severe consequences, including ecosystem contamination and a decline in biodiversity. Thus, the adoption of sustainable agricultural strategies has gained traction over the years, including the use of microbial inoculants containing Plant Growth-Promoting Bacteria (PGPB) (Shahwar *et al*., 2023). These microorganisms promote plant growth through various mechanisms, including biofertilization, production of growth-regulating hormones, countering pathogens and pests, and enhancing plant stress tolerance (Kumar *et al*., 2022; Bittencourt *et al*., 2023).

Central to this field is *Azospirillum*, a genus of Gram-negative, diazotrophic Alphaproteobacteria. First identified in 1925 and gaining prominence in the 1970s (Cassán *et al*., 2020), these microbes are known for their metabolic versatility and wide distribution (Cassán *et al*., 2020); species of *Azospirillum* are commonly found in the root zones of critical crops such as maize and rice, and have been discovered in a vast range of environments, from high-salinity soils and peat bogs (Reinhold *et al*., 1987; Doroshenko *et al*., 2007) to contaminated industrial sites (Ojeda-Morales *et al*., 2015), demonstrating the genus exceptional adaptability and potential.

A combination of direct and indirect plant-growth-promoting mechanisms drives the beneficial effects of *Azospirillum*. While Biological Nitrogen Fixation (BNF) was the first major trait studied in *Azospirillum*, the production of phytohormones, particularly Indole-3-Acetic Acid (IAA), is now recognized as the key mechanism underlying its beneficial effects, primarily through the enhancement of root system architecture (Bashan and Levanony, 1990). Furthermore, these bacteria employ a suite of other strategies, including phosphate solubilization, production of iron-scavenging siderophores, and synthesis of various other growth regulators that work together to boost plant health and resilience (Bottini *et al*., 1989; Cohen *et al*., 2009; Kusajima *et al*., 2018).

The agricultural significance of *Azospirillum* is underscored by its extensive use in commercial inoculants, especially within the large-scale farming systems in South America (Cassán *et al*., 2020). These products play a vital role in moving towards more sustainable practices by reducing the need for synthetic fertilizers. However, the current market is surprisingly limited in taxonomic diversity. The majority of the available inoculants are formulated using just two closely related species: *A. brasilense* and *A. argentinense*, with a single strain, Az39, dominating commercial products (Cassán *et al*., 2020). This reveals a remarkable contrast between the genus diversity (which has a total of 31 species identified in NCBI databases) and its phylogenetically narrow application in agriculture.

While the capabilities of important agricultural strains of *Azospirillum* have been revealed through sequencing and genomic inference, commercial interest has largely remained focused on nitrogen fixation. A broader, systematic genomic exploration of the entire genus would open opportunities to investigate novel strains and uncover additional mechanisms of plant growth promotion. In this work, we report a comprehensive comparative genomic analysis of publicly available *Azospirillum* genomes, describing their pangenome and genetic repertoire. In addition, we sequenced and analyzed four novel *Azospirillum* isolates obtained from a large local collection of microorganisms of biotechnological interest. These strains isolated from plant sources were selected based on their potential as candidates for bioconsortia. Our analyses reveal an *Azospirillum* species cluster containing species commonly associated with plant growth promotion. This cluster presents a large set of exclusive genes, mainly linked to chemotactic capability, which appear to define the cluster’s lifestyle. Together, these findings point to the existence of novel species with plant growth-promoting potential, broadening biotechnological applications of the genus *Azospirillum*.

## Methods

### Genome sequencing

Four bacterial isolates collected between August and October 2022 were selected from the Collection of Microorganisms of Biotechnological Interest at LBCT-UENF (Table 1). Before genome sequencing, isolates were cultivated in Nutrient Broth (NB) medium to ensure viability and growth. Preliminary identification through 16S rRNA gene sequencing was conducted in a sequencing facility (NGS - Piracicaba, Brazil). Bacterial DNA samples were extracted using the Wizard Genomic DNA Purification kit (Promega - Madison, WI, USA) and sequenced on the PromethION 2 Solo system at the Institute of Biodiversity and Sustainability (UFRJ, Macaé, Brazil). Sequencing quality was assessed using FastQC v0.12.0. Low-quality reads were trimmed using Trimmomatic v0.39 (Bolger, Lohse and Usadel, 2014), and genomes were assembled with SPAdes v4.2.0 (Prjibelski *et al*., 2020).

**Table 1:** Newly sequenced *Azospirillum* isolates from the UENF collection and their sources of isolation.

| Name | Proposed name | Source | 16S identification | GenBank accession |
| --- | --- | --- | --- | --- |
| New isolate A | *Azospirillum* sp. UENF 112521 | Pineapple roots | *Azospirillum* sp. | Submitted to GenBank* |
| New isolate B | *Azospirillum* sp. UENF 111221 | Pineapple rhizosphere | *Azospirillum palustre* | Submitted to GenBank* |
| New isolate C | *A. brasilense* UENF 312511 | Papaya roots | *Azospirillum formosense* | Submitted to GenBank* |
| New isolate D | *A. argentinense* UENF 211521 | Guava rhizosphere | *Azospirillum formosense* | Submitted to GenBank* |

\* Awaiting response

### Collection and curation of publicly available genomes

A total of 296 Azospirillaceae genomes were downloaded from the GenBank database using NCBI Datasets v2 (https://www.ncbi.nlm.nih.gov/datasets/) in July 2024 (Table S1). Genome quality was assessed using BUSCO v5.8.2 (Seppey, Manni and Zdobnov, 2019). Genomes exhibiting less than 95% completeness, more than 5% duplication, or more than 500 contigs were discarded. Pairwise genomic identity was estimated with Mash v2.3 (Ondov *et al*., 2016) to estimate pairwise genomic identity (1 - Mash distance). Also, genomes showing more than 99% identity were considered redundant; in such cases, the assembly with superior quality metrics (contig number, L50, and N50) was retained for downstream analyses.

### Genus delimitation and phylogenetic analysis

To resolve the structure of the *Azospirillum* genus within our dataset, we performed Average Nucleotide Identity (ANI) analysis (PyAni v0.3.0-alpha). ANI values were calculated for the 119 curated genomes (115 public genomes, plus our four newly sequenced isolates). *Rhodospirillum rubrum* (GCA_000013085.1) was included as an outgroup to root the phylogeny. Genomes forming a coherent cluster identified as *Azospirillum* in the ANI analysis were retained for downstream analyses (Table S1).

Additionally, *Azospirillum* groups were delineated using a graph-based approach derived from Mash distances, as previously described (Passarelli-Araujo *et al*., 2021) and (Henaut-Jacobs, Passarelli-Araujo and Venancio, 2023). For this analysis, we used the genomes from the *Azospirillum* cluster identified in the ANI step. Mash distances (1−Mash values) were used to estimate pairwise genomic relationships, with genomes represented as vertices in the resulting graph.

A genus-level phylogeny was also inferred from a species tree constructed with orthologous genes identified by Orthofinder v3.0.1 b1 (Emms and Kelly, 2019). A maximum-likelihood phylogenetic tree was then generated using IQTree2 v2.3.6 (Minh *et al*., 2020), with *R. rubrum* serving as the outgroup.

### Pangenome analysis

*Azospirillum* genomes were annotated with Prokka v1.14.6 (Seemann, 2014). Pangenome analysis was then carried out using Roary v3.13.0 (Page *et al*., 2015) with an 80% identity threshold for orthology assignment. Genome clustering was guided by the original genus phylogeny established in the previous session. The pangenome was partitioned into four categories based on gene prevalence across genomes: Core (>99%), Soft core (99%-95%), Shell (95%-15%), and Cloud (<15%) (Matthews *et al*., 2024).

### Accessory pangenome functionality

After analyzing the clustering of the accessory pangenome across *Azospirillum* genomes, we selected the cluster containing the most agriculturally relevant genomes, which formed a single community (See “*A. brasilense* is part of a great *Azospirillum* species complex” in the Results). A reference genome for this community (GCA_022023855.1) was chosen based on assembly quality metrics (completeness and N50). This reference genome was submitted to STRING for functional annotation of exclusive genes in the complex. Proteins annotated as hypothetical or unknown were manually curated and renamed using UniProt information. Protein-protein interactions were further explored in Cytoscape.

Genes exclusive to the *A. brasilense* species complex were also screened for their orthologs hosted in STRING (Szklarczyk *et al*., 2025). The corresponding accession codes were retrieved using the STRING plugin of Cytoscape v3.10.2 (Kohl, Wiese and Warscheid, 2011), applying a confidence score threshold of 0.9. Singlets and clusters containing fewer than three genes were discarded. The remaining genes were reannotated with InterProScan (Jones *et al*., 2014).

### Specific traits profiling

Given the substantial agricultural relevance of the *Azospirillum* genus, we investigated key traits related to antibiotic resistance and plant growth-promotion potential. Antibiotic resistance profiles were inferred for all genomes using The Comprehensive Antibiotic Resistance Database (CARD) v3.1.3 (Alcock *et al*., 2023), a reference database for antibiotic resistance-associated genes. Usearch v11.0.667 (Zhou, Liu and Li, 2024) was employed to align *Azospirillum* genomes to CARD with a 60% identity cutoff. Simultaneously, plant growth-promotion potential was assessed by screening genomes against an in-house curated database, also utilizing Usearch. This database comprises genes associated with direct plant-growth promotion, such as those involved in nitrogen fixation, phosphate solubilization, and phytohormone biosynthesis. To reduce phylogenetic bias, we incorporated diverse reference sequences for each gene, ensuring robust inference of gene presence regardless of evolutionary distance from the reference sources.

## Results and Discussion

### The *Azospirillum* genus is well established within the Azospirilaceae family

ANI analysis and graph-based Mash distance clustering allowed us to precisely define the boundaries of the *Azospirillum* genus within our dataset. The vast majority of genomes previously classified as *Azospirillum* formed a dense and cohesive cluster, while a small subset had weak relationships with this core group and were excluded from the downstream analyses due to their affiliation with closely related taxa (Figure 1). The genomes of our four newly sequenced isolates exhibited a strong relationship with other *Azospirillum* genomes, corroborating their preliminary 16S rRNA-based classification (Table 1, Figure S1).

**Figure 1.**
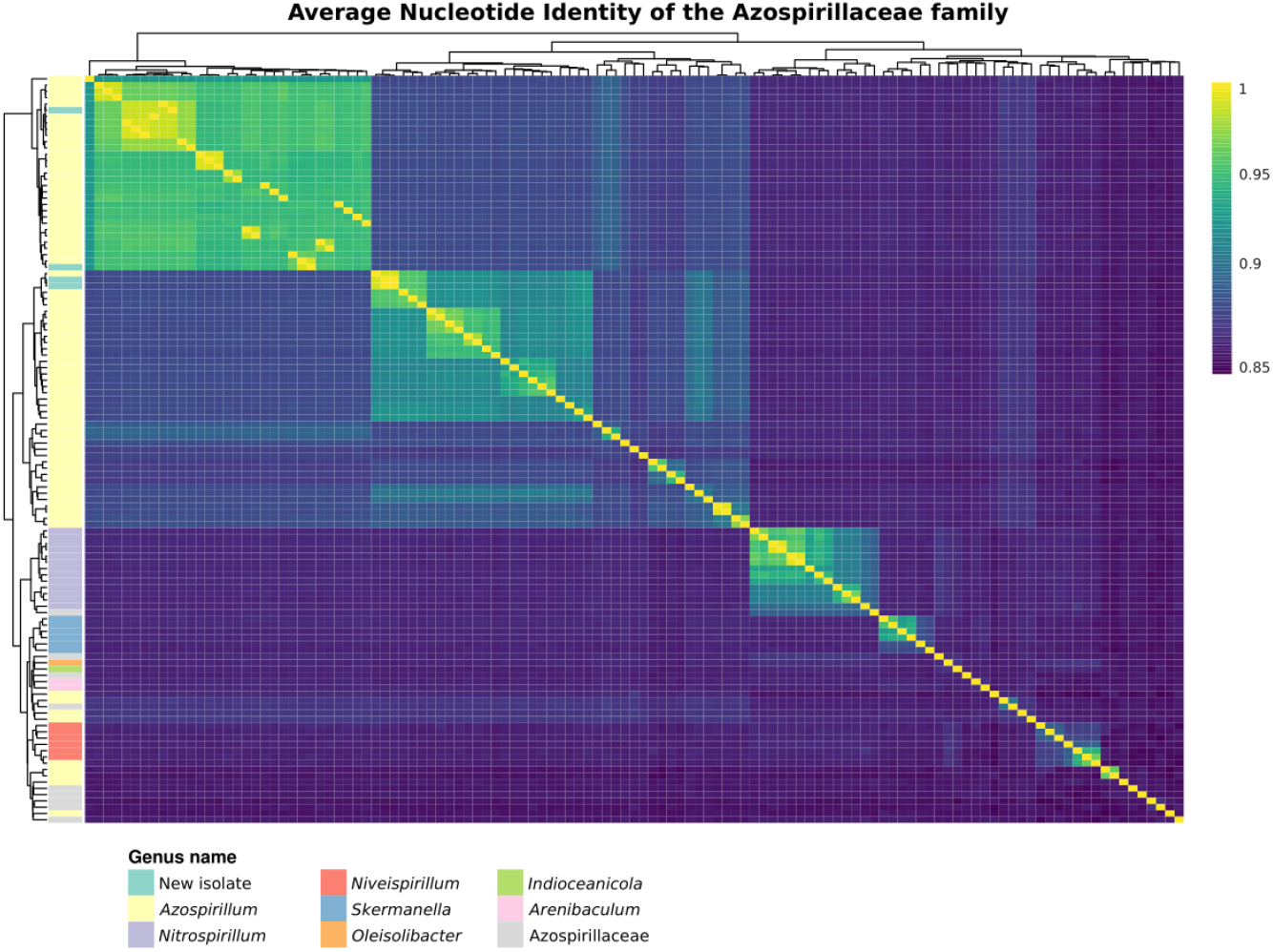
Heatmap using ANI values between every pair of genomes (represented as 1-Mash). Colors in the left annotation bar represent species classification in GenBank.

Genomic clustering further revealed two major groups within the genus. One of these clusters notably encompassed numerous genomes with well-established plant growth-promoting capacities, particularly *A. brasilense* and *A. argentinense*. This clear pattern of relatedness, highlighting agriculturally relevant species, provided the framework for our subsequent in-depth analyses of *Azospirillum* taxonomic classification and functional potential.

### *A. brasilense* is part of a greater *Azospirillum* species complex

The precise definition of bacterial species remains a challenge in microbial ecology, often debated as to whether genetic diversity exists on a continuum or in discrete groups (Caro-Quintero and Konstantinidis, 2012; Cohan, 2019). Genetic discontinuity, representing abrupt breaks in genomic identity, is a key concept for delineating species boundaries. While a 95% ANI is a widely accepted threshold for species classification (Konstantinidis, Ramette and Tiedje, 2006; Barghouthi, 2011), recent research quantifies this discontinuity, termed δ (Passarelli-Araujo, Venancio and Hanage, 2025), by measuring the steepest change in genomic identity within a genomic dataset. Further, this quantitative approach reveals a significant (Brockhurst *et al*., 2019; Dewar *et al*., 2024) association between genetic discontinuity and a species’ lifestyle, primarily reflected in its pangenome features. Species with high genetic discontinuity often exhibit closed pangenomes, suggesting specialized lifestyles with limited gene exchange (Hollensteiner *et al*., 2023; Dewar *et al*., 2024). Conversely, species with more open pangenomes and versatile environmental adaptations tend to show less pronounced, yet still discernible, genetic breaks (Brockhurst *et al*., 2019). This spectrum of discontinuity, where clear divisions are sometimes evident and at other times ambiguous, parallels the challenges encountered in defining “ring species” in ecology, where gradual changes across populations eventually lead to distinct forms without clear points of separation (Alcaide *et al*., 2014). Understanding these varying degrees of genetic discontinuity is vital for reassessing traditional bacterial species classifications and appreciating the fluidity of microbial evolution.

In our investigation of *Azospirillum* species delineation, we detected a notable species complex within broader patterns of genetic discontinuity. Most well-characterized species, particularly those already used in commercial inoculants (Cassán and Diaz-Zorita, 2016; Dos Santos Ferreira *et al*., 2022; Maroniche *et al*., 2024), were found to be closely related, forming a concise cluster. The canonical 95% ANI threshold proved too stringent, often fragmenting well-supported taxa (Figure 2). Reducing the threshold slightly to 94% effectively uncovered a cohesive species complex that encompasses *A. brasilense, A. argentinense, A. baldaniorum, A. formosense, A. aestuarii*, and *A. tabaci* (Figure 2B). Importantly, two of our novel isolates (C and D) were also placed within this complex. We hereafter refer to this group as the *A. brasilense* complex. Using this 94% identity cutoff, we identified a total of 25 genomic communities (Table S2).

**Figure 2.**
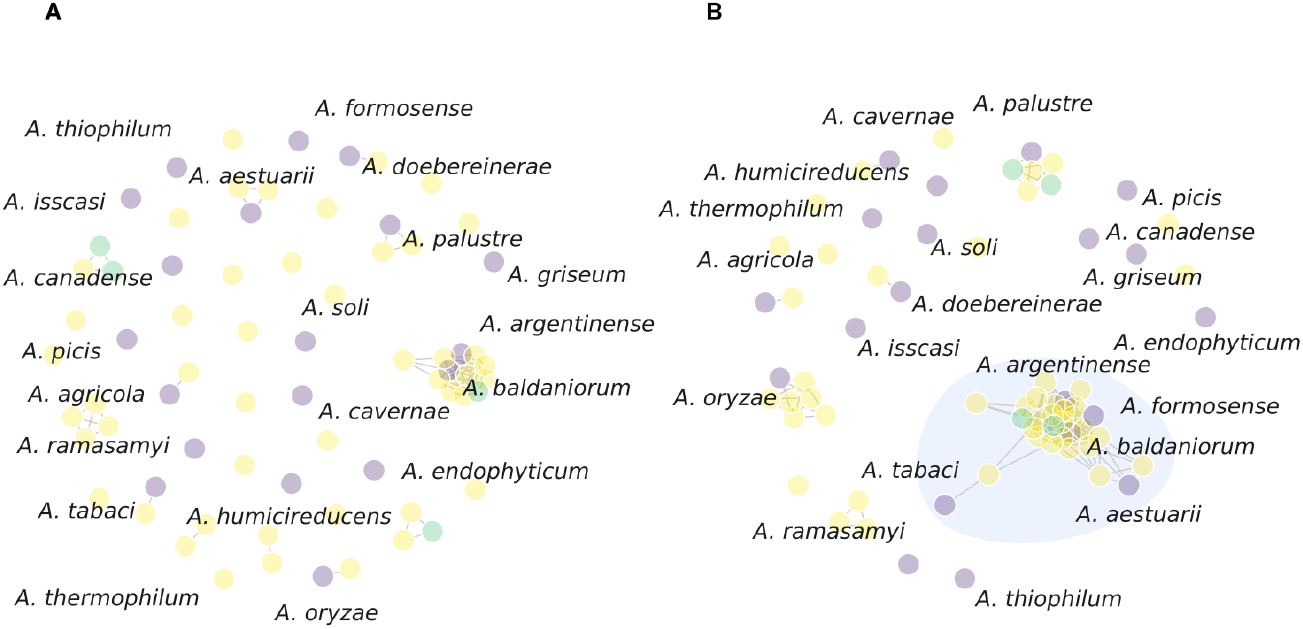
Graph-based approach to species delineation. Purple vertices represent type genomes that passed through quality and redundancy analysis, green vertices represent the new *Azospirillum* isolates introduced in this study, and yellow vertices represent all other genomes. A) Clustering using a 95% identity threshold; B) Clustering using a 94% identity threshold, the highlighted cluster is the *A. brasilense* species complex.

Phylogenetic analysis based on the species tree of orthologous genes confirmed the strong genetic relationships within the *A. brasilense* complex (Figure 3). This tree also reflected the broader clustering patterns observed in our genus-wide classification, with a clear separation of the *A. brasilense* complex. Beyond this group, the lower cluster of the tree included additional *Azospirillum* groups. In contrast, the other large cluster consisted of multiple independent species, comprising twelve previously categorized reference genomes and 16 uncharacterized strains.

**Figure 3.**
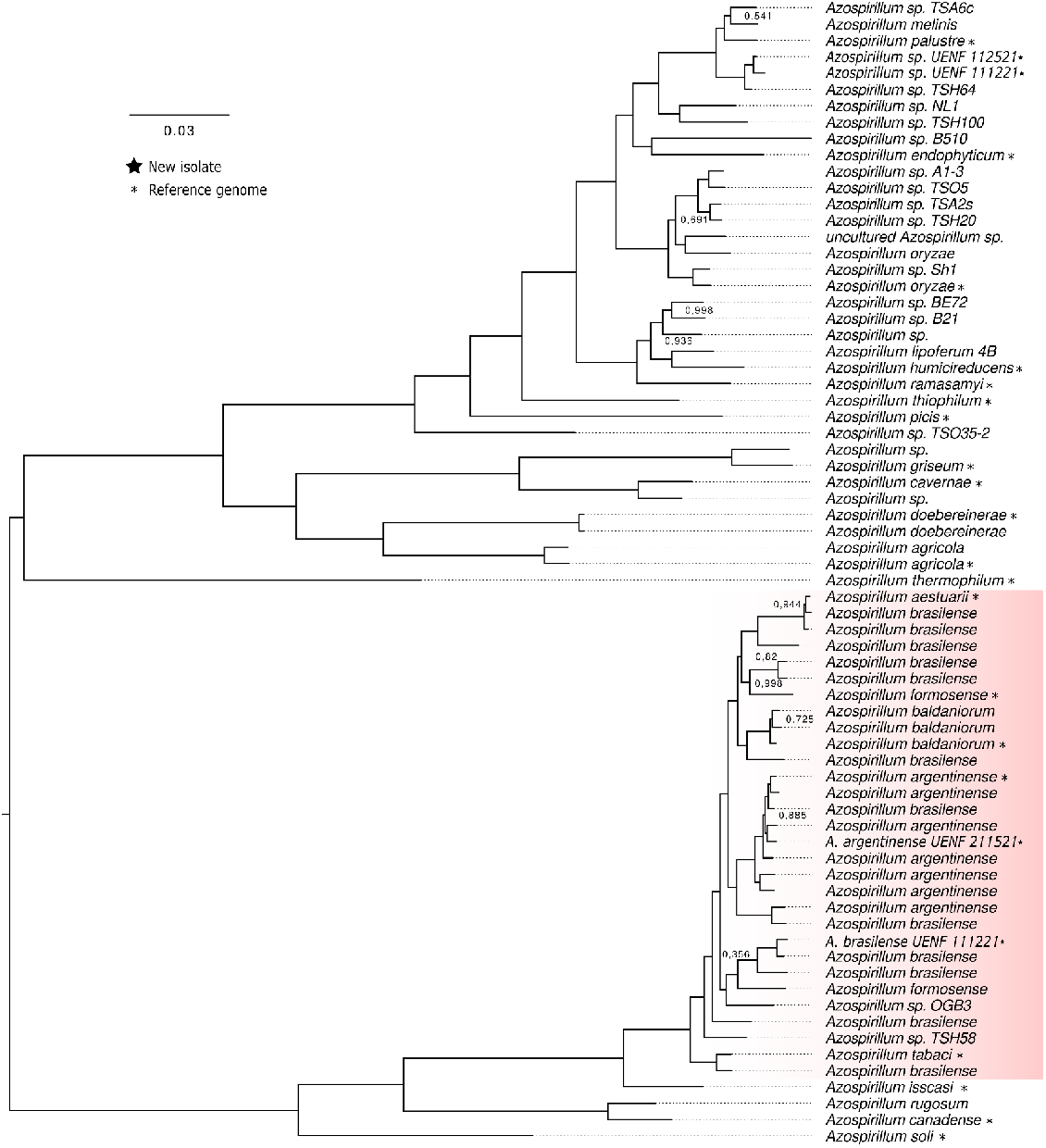
Phylogenetic tree based on single-copy orthologs from each genome. The red cluster represents the *A. brasilense* species complex defined in this study. Bootstrap values of 1 were omitted for clarity.

Despite the cladogram topology within the *A. brasilense* complex suggesting that these genomes form a single species, evidence from the literature indicates otherwise. Phenotypic differentiation among strains—such as variation in preferred pH and carbon sources (e.g., D-fructose, maltose)—supports their recognition as distinct species (Xu *et al*., 2023). Nonetheless, these same taxa exhibit very limited genomic differentiation, as revealed by multiple clustering approaches, including both ANI values and phylogenetic analyses of orthologous genes.

This phylogenetic classification also allowed us to infer the taxonomic position of *A. brasilense UENF 111221* and *A. argentinense UENF 211521*, both of which were placed in small, distinct monophyletic clades.

### Pangenome analysis revealed an exclusive gene cluster in the *A. brasilense* species complex

At first glance, our dataset revealed two distinct clades of closely related genomes. Despite this high genetic similarity, the scientific literature consistently classifies these genomes as separate species. To further explore this separation, we conducted a super-pangenome analysis to identify the genetic determinants underlying the division between the two clades, with particular emphasis on the gene sets associated with the agriculturally important *A. brasilense* species complex.

Our analysis showed that the *Azospirillum* genus harbors a large and diverse pangenome, characterized by a very small set of conserved core genes (Figure 4). In total, we identified 866 core, 320 soft-core, and 9,163 shell gene families. The vast majority of the repertoire consisted of 56,921 cloud genes, representing 84.6% of the total pangenome and highlighting the remarkable genome plasticity of the genus (Figure S2). This plasticity is likely shaped by frequent horizontal gene transfer within the complex polymicrobial environments that *Azospirillum* species inhabit (Georgiades and Raoult, 2010). Notably, the shell pangenome (orthologs present in 15–95% of genomes) exhibited a distinct structure, with gene families exclusive to one of the major communities but absent in the other.

**Figure 4.**
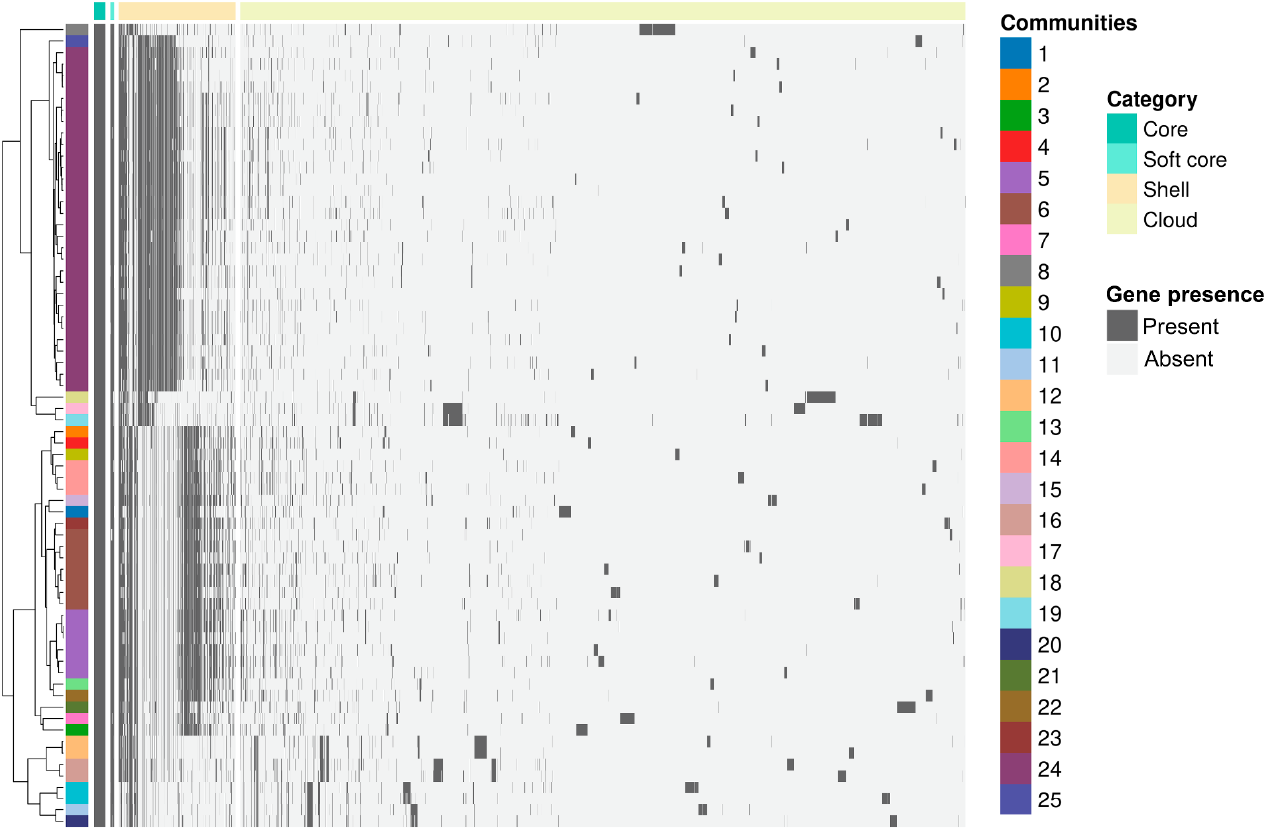
Pangenome of the *Azospirillum* genus. The top top annotation bar indicates the distribution of core, soft-core, shell, and cloud genes. The left annotation bar identifies and labels communities containing more than one genome.

To further investigate the genomic basis for the phylogenetic groupings, we searched for clade-specific markers. One clade, in particular, stood out: the *A. brasilense* species complex (community 24) together with the species corresponding to communities 17 (*A. rugosum*), 18 (*A. soli*), and 19 (*A. canadense*). This group alone contained 2,471 orthologous gene clusters unique to its members. Although many of these orthogroups were annotated as hypothetical proteins (Table S3), we leveraged the STRING database to predict their putative functions and gene interaction networks based on sequence similarity (Figure 5).

**Figure 5.**
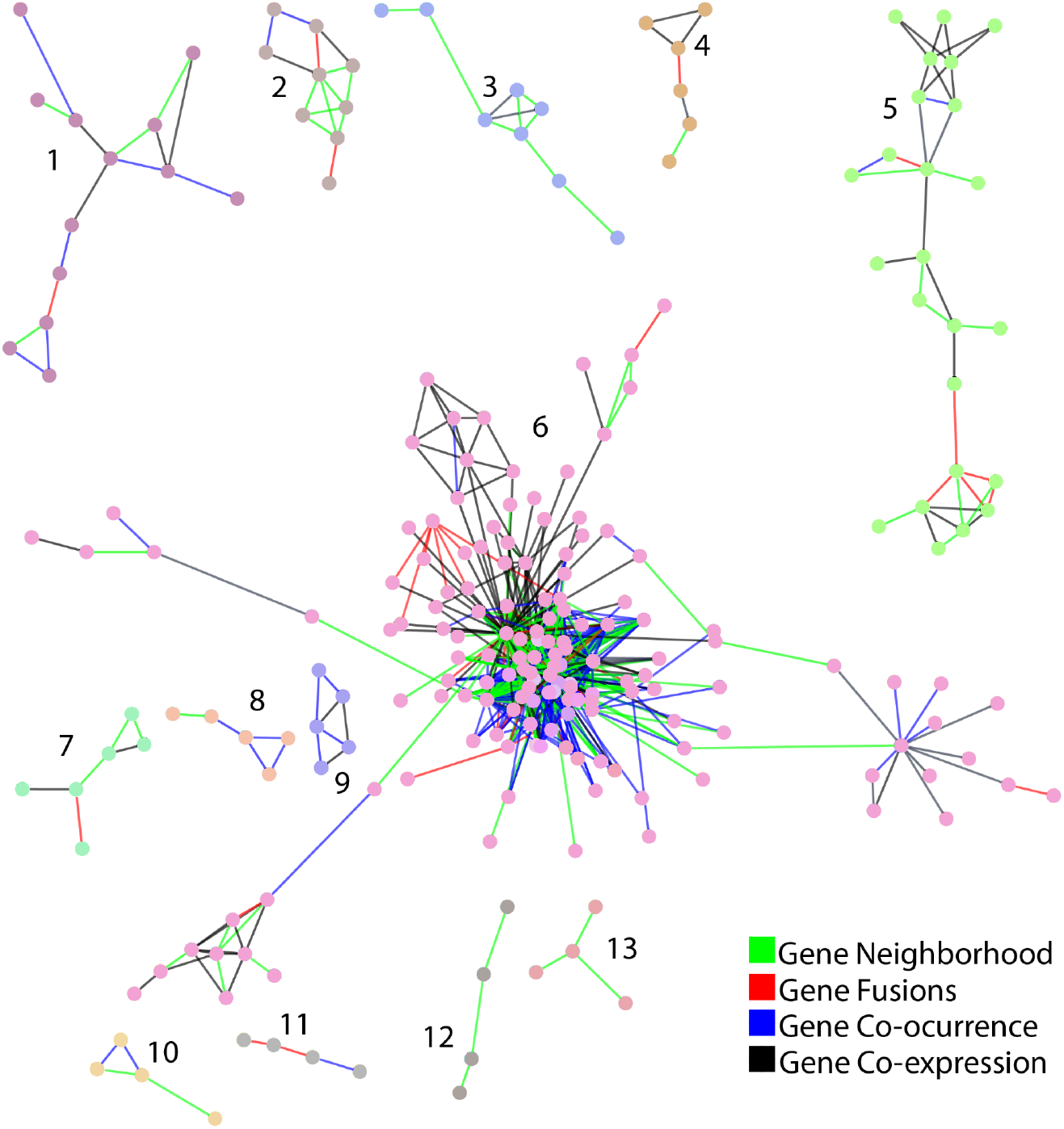
STRING network of the genes exclusive to the *A. brasilense* complex. Cluster content for each numbered group is detailed in Table S4. Edge colors indicate gene relationships (neighborhood, fusion, co-expression, or co-occurrence) with a confidence score >0.9. Node colors represent distinct clusters. Distances are not to scale.

The STRING network of genes exclusive to the *A. brasilense* species complex revealed 13 clusters, each containing at least four genes. Functional annotation indicated a high abundance of duplicated and functionally related genes, including ABC transporters (G3DSA:3.40.50.300:FF:000421, IPR022467, IPR022478, IPR051449, IPR050319, IPR050388, and IPR051120), adenylyl/guanylyl (IPR050697) and diguanylate cyclases (IPR050469), chemotaxis methyl-accepting receptors (IPR004090), and at least 37 genes associated with histidine kinases and their regulation, particularly protein families PF02518, IPR050736, IPR050980, and G3DSA:3.30.565.10:FF:000010. Full details for each cluster are provided in Table S4 and Figure S3.

Our findings highlight a highly specialized gene repertoire that is strongly linked to chemotaxis and plant-host interactions, including motility and the ability to sense environmental gradients. We hypothesize that this genomic toolkit confers an evolutionary advantage to the *A. brasilense* species complex, which may underlie its success as a commercial bioinput and enhance the potential use of other species within the same cluster.

### Screening for genes associated with plant growth-promotion shows their uniform presence within the genus

We assessed the plant growth-promotion (PGP) potential of all *Azospirillum* genomes using an in-house, literature-based reference dataset. This analysis revealed a conserved core set of PGP genes across the genus, including those involved in nitrogen fixation and phosphate solubilization (Figure 6). Notably, the *A. brasilense* complex harbored the *ipdC* gene, a central gene for the production of indoles, primarily IAA, an auxin that regulates root development, nutrient uptake, and overall plant growth (Sun *et al*., 2022). This could be linked to the usage of already known PGP species from this complex, and highlights the untapped potential of less explored members, such as *A. tabaci* and *A. formosense*, both soil isolates with documented PGP capacity (Lin *et al*., 2012; Duan *et al*., 2021) but not yet commercialised in bioconsortia. Importantly, our novel isolates C and D also share this PGP potential with species already recognized for their beneficial interactions with plants.

**Figure 6.**
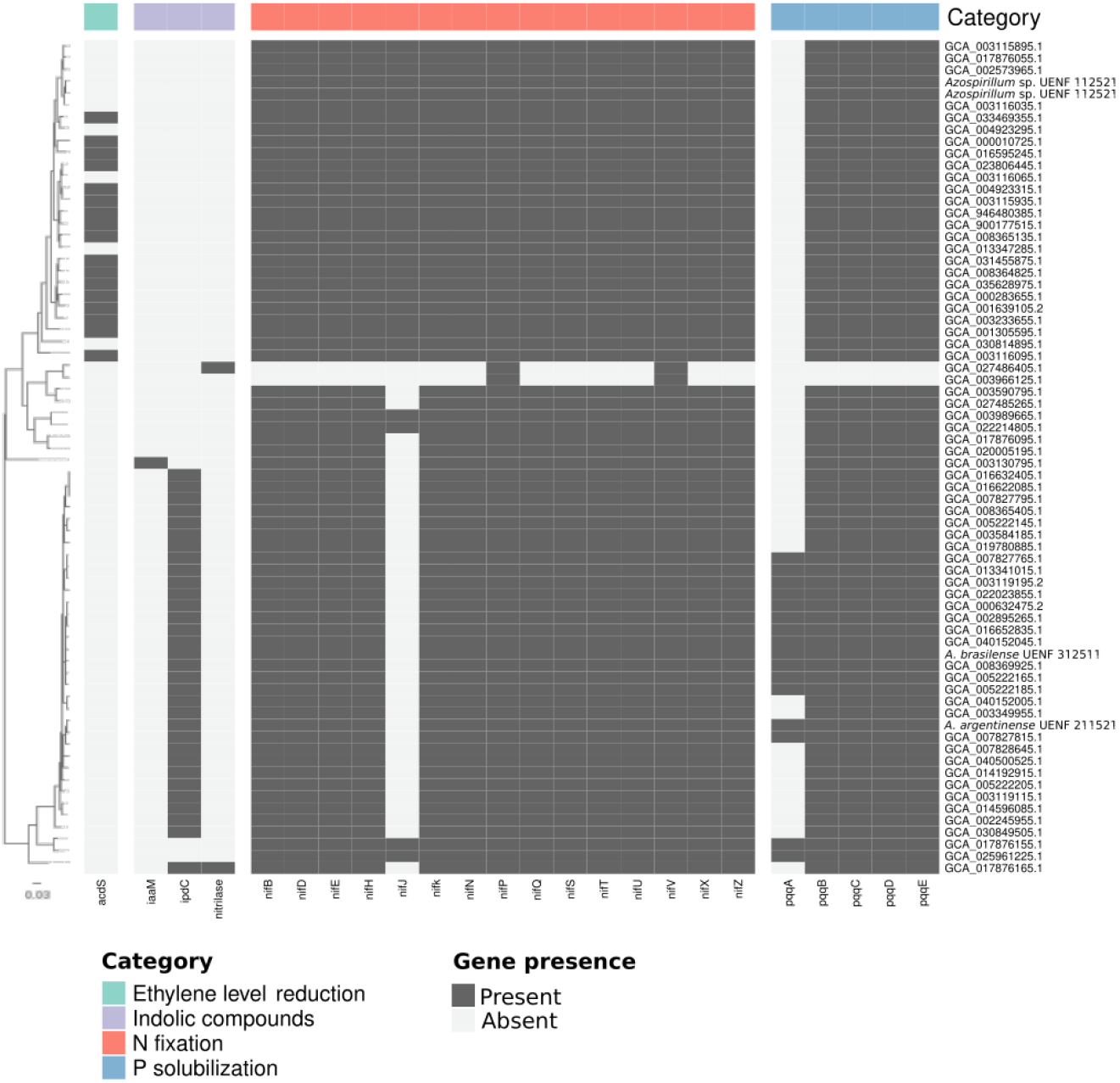
Presence/absence heatmap of direct plant growth-promoting genes. The top annotation bar indicates the main functional categories associated with each gene.

Furthermore, a subset of the *A. brasilense* species complex also carried the *pqqA* gene, which was absent from the rest of the genus (Figure 6). *pqqA* is important for the biosynthesis of pyrroloquinoline quinone (PQQ), a coenzyme that plays a key role in various metabolic processes, such as the oxidation of sugars and alcohols (Kim *et al*., 2003). Nonetheless, previous studies have demonstrated that the PQQ operon can function without *pqqA* (Matteoli *et al*., 2018). In the context of phosphate solubilization, PQQ-dependent enzymes—particularly glucose dehydrogenase—play a crucial role in releasing insoluble phosphate from the soil, thereby enhancing plant nutrient acquisition.

Although PGP genes are broadly conserved across *Azospirillum*, two genomes showed a markedly reduced repertoire: *Azospirillum* sp. C1_MAG_00050 (GCA_027486405.1 (unpublished)) and *Azospirillum griseum* L-25-5 w-1 (GCA_003966125.1 (Yang *et al*., 2019)). Both genomes were isolated from lake water samples, with the former being a metagenome-assembled genome (MAG). Their limited PGP gene content may reflect ecological adaptation, as aquatic environments likely impose weaker selective pressure for plant-associated traits.

We also analyzed antibiotic resistance genes to infer the biosafety profile of the *Azospirillum* genomes. No concerning resistance traits were detected in any of the genomes examined. The unique presence of specific plant growth promotion genes (*ipdC, pqqA*) in the *A. brasilense* species complex (Figure 7), together with this favorable biosafety profile, underscores the potential of these strains for the safe development and application of biofertilizers.

**Figure 7.**
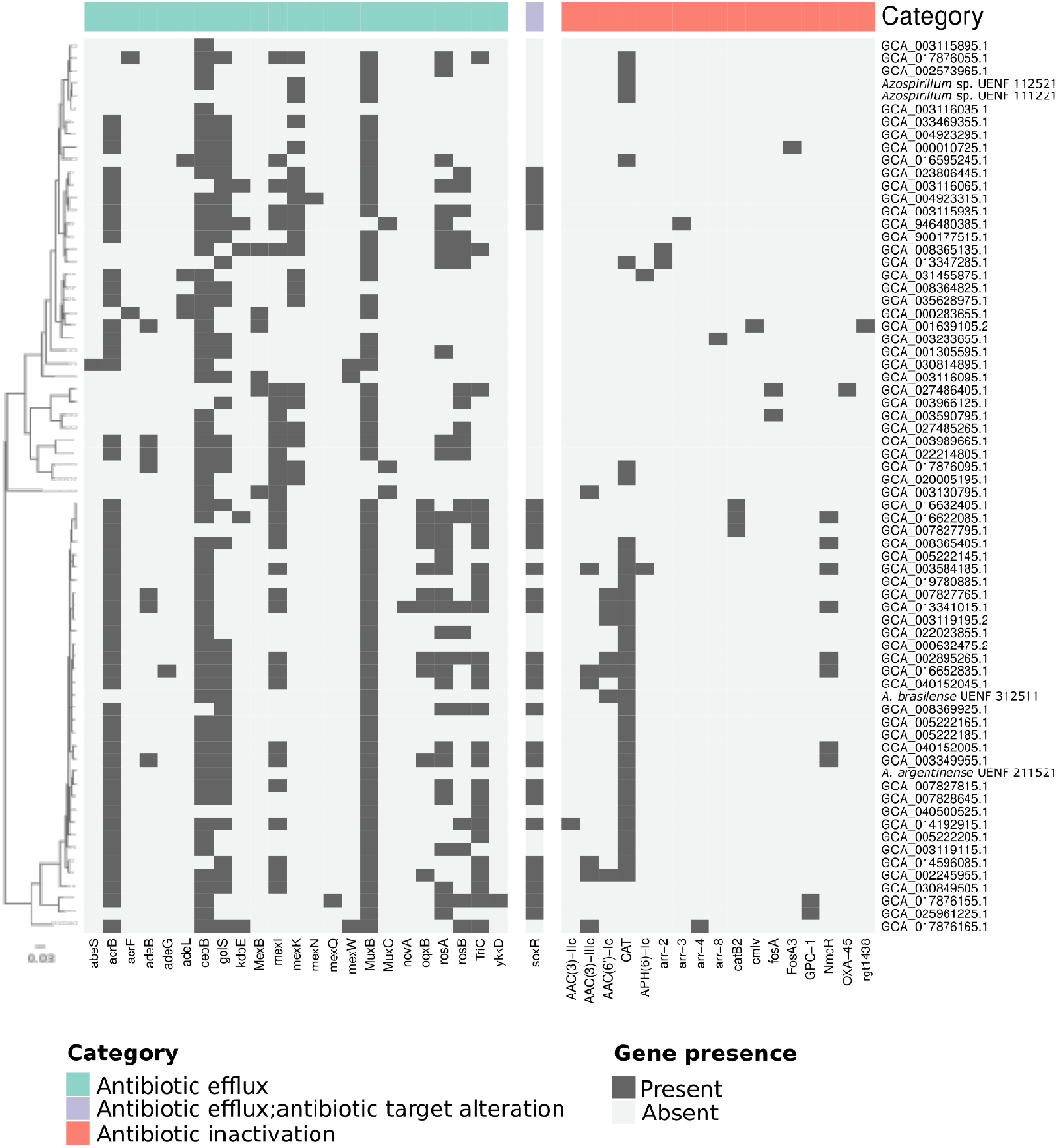
Presence/absence heatmap of antibiotic resistance genes. The top annotation bar indicates the presumed mode of action for each gene.

Considering the collective evidence, we propose that the species clustered within the *A. brasilense* species complex represent up-and-coming candidates for microbial inoculant applications. This conclusion is supported by their consistently favorable biosafety profiles and their shared genetic repertoire of key plant growth-promoting traits. This research expands the scope of potential bioinputs by highlighting less-explored *Azospirillum* species such as *A. formosense* and *A. baldaniorum*. Although *A. baldaniorum* was only recently described (2020), it already shows a promising genomic toolkit for phytohormone production (Dos Santos Ferreira *et al*., 2020). Similarly, *A. formosense* has demonstrated the ability to engage in synergistic interactions with other beneficial microorganisms, such as *Bacillus* species (Yaadesh *et al*., 2023), but it remains underutilized in agricultural applications. By characterizing these species and reporting novel isolates within this complex, our study underscores their potential as next-generation biofertilizers.

## Conclusion

This comprehensive genomic study of *Azospirillum* refines current understanding of species boundaries, identifying a major species complex characterized by high genomic identity despite phenotypic distinctiveness. Pangenome analysis revealed unique genetic features within this complex, notably the presence of *ipdC* and *pqqA* genes, both central to plant growth promotion. Combined with a favorable biosafety profile and the distinct ecological and physiological traits of the newly defined *A. brasilense* species complex, these findings underscore the considerable yet underexplored biotechnological potential of *Azospirillum* species beyond those currently in use. Our work paves the way for the development of more effective and diverse microbial inoculants to advance sustainable agriculture.

## Acknowledgements

This work was supported by Fundação Carlos Chagas Filho de Amparo à Pesquisa do Estado do Rio de Janeiro (FAPERJ E-26/210.291/2021), Coordenação de Aperfeiçoamento de Pessoal de Nível Superior-Brasil (CAPES; Finance Code 001), Conselho Nacional de Desenvolvimento Científico e Tecnológico (CNPq) and Programa de Apoio à Pesquisa, Inovação e Cultura (PAPIC - UENF). The funding agencies had no role in the design of the study, the collection, analysis, and interpretation of data, or the writing. The authors would like to thank Dr. Vitor B. Pinto for his valuable assistance with the PPi network configuration.

## Author contributions

Conceptualization: S.H.-J., I.O.P, and T.M.V.; formal analysis: S.H.-J.; computational methodology: S.H.-J., F.F.R.-C, and G.Q.-P.; wet lab methodology; S.H.-J., G.P.-M., and B.C.R.; writing—original draft: S.H.-J. and T.M.V.; writing—review and editing: S.H.-J., R.S.E. and T.M.V.; funding acquisition: R.N.-F., F.L.O. and T.M.V.; project administration: F.L.O. and T.M.V. All authors have read and agreed to the published version of the manuscript.

