## Supplementary figures for "Genomic plasticity of the *Azospirillum* genus in a biotechnological context"

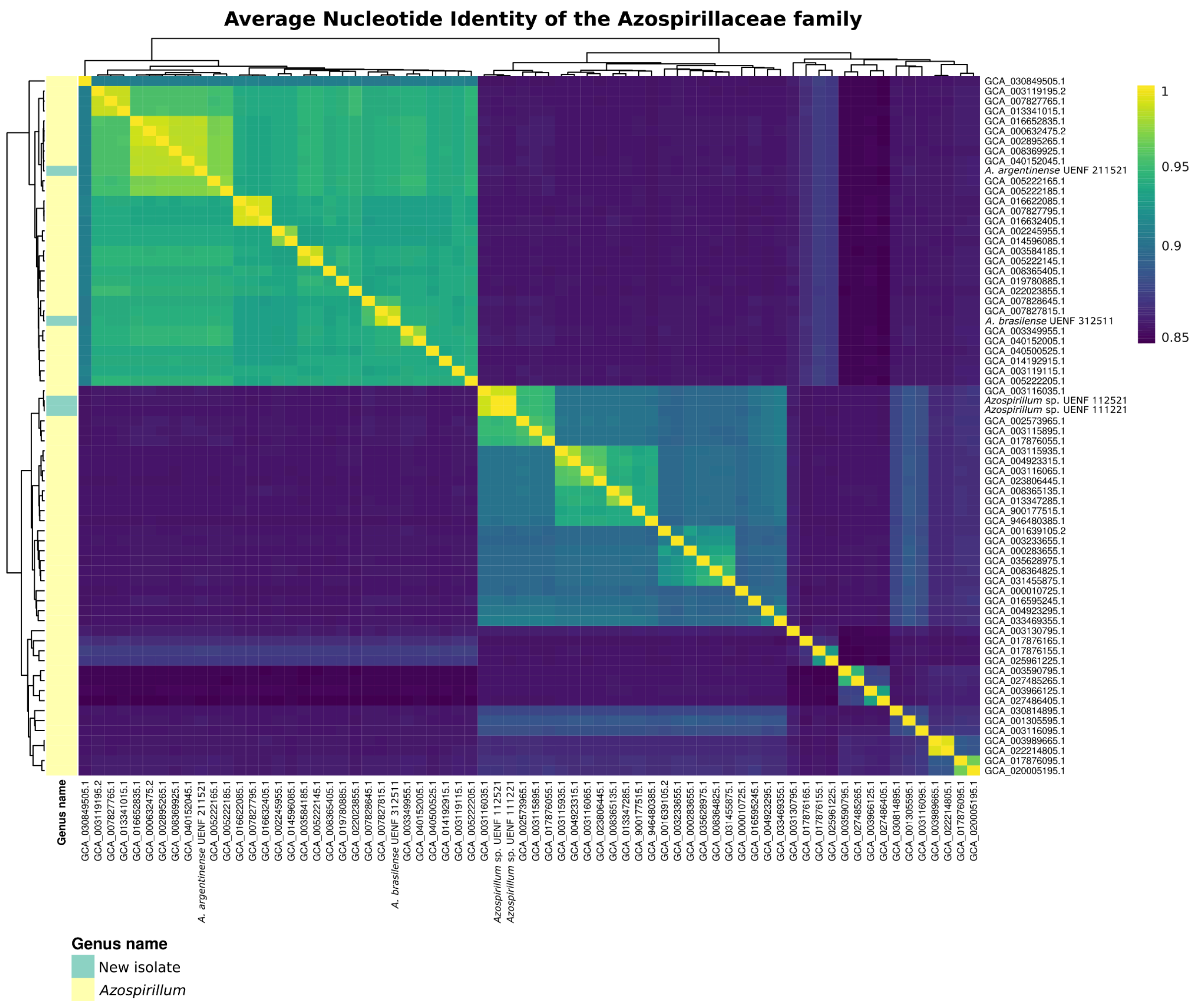


Figure S1. The final genomes of *Azospirillum* heatmap using ANI values between every pair of genomes. Colors in the left annotation bar represent species classification in GenBank.

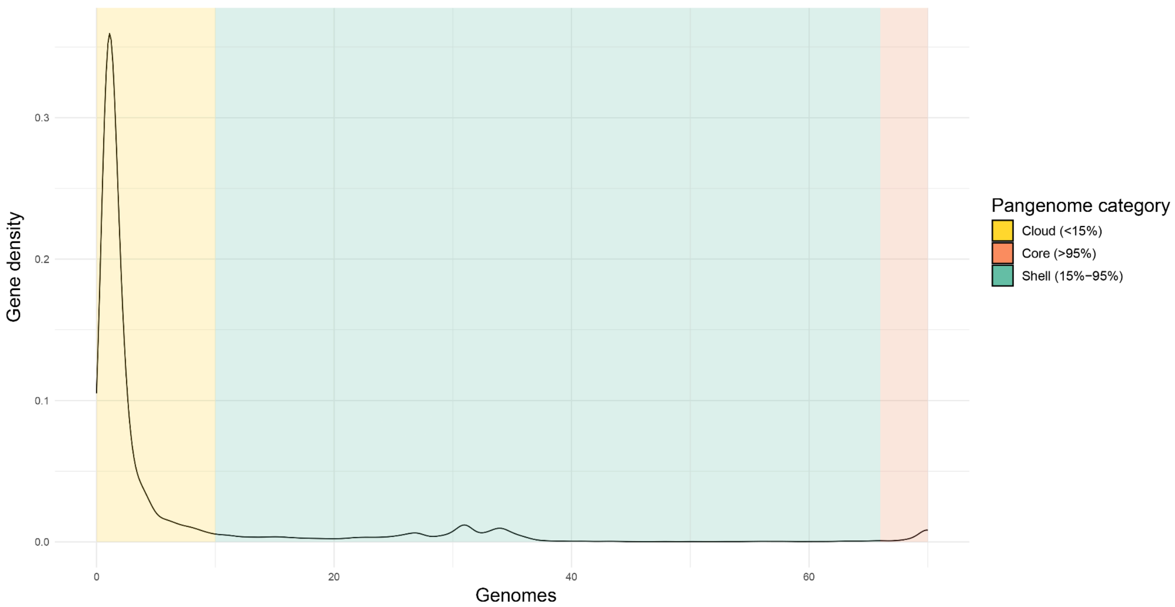


Figure S2: Distribution of genes across *Azospirillum* genomes. The pangenome is divided into three categories: Cloud genes, comprising rare and strain-specific genes; Shell genes, present in subsets of genomes and often associated with specific clusters; and Core genes, conserved across nearly all genomes and essential to the genus.

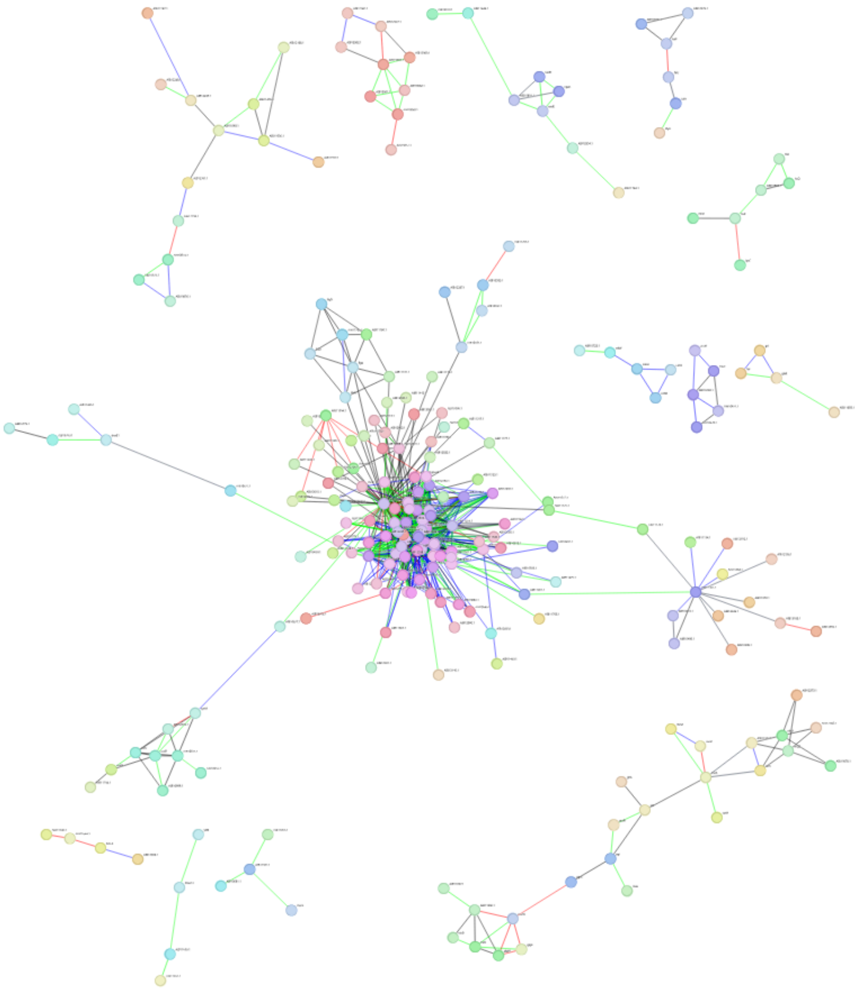

Figure S3: STRING network of the *A. brasilense* complex-exclusive genes with labels. The cluster content according to each number is detailed in Table S4. Edge (line) color indicates gene relationship (neighborhood, fusion, co-expression, or co-occurrence) with >0.9 confidence score. Different vertex colors represent different clusters. Distances do not represent real scales.
